# Comparative evaluation of four exome enrichment solutions in 2024: Agilent, Roche, Vazyme and Nanodigmbio

**DOI:** 10.1101/2024.07.11.602872

**Authors:** Vera Belova, Iuliia Vasiliadis, Zhanna Repinskaia, Alina Samitova, Anna Shmitko, Natalya Ponikarovskaya, Oleg Suchalko, Valery Cheranev, Shatalov Peter, Shegai Peter, Kaprin Andrey, Denis Rebrikov, Dmitriy Korostin

## Abstract

Whole exome sequencing (WES) is essential for identifying genetic variants linked to diseases. This study compares available to date four exome enrichment kits: Agilent SureSelect Human All Exon v8, Roche KAPA HyperExome Probes, Vazyme VAHTS Target Capture Core Exome Panel, and Nanodigmbio NEXome Plus Panel v1. We evaluated target design, coverage statistics, and variant calling accuracy across these four different exome capture products. All kits showed high coverage completeness, with mean x10 coverage exceeding 97.5% and x20 coverage above 95%. Roche exhibited the most uniform coverage, indicated by the lowest fold-80 scores, while Nanodigmbio had more on-target reads due to fewer off-target reads. Variant calling performance, evaluated using in-lab standard E701 DNA sample, showed high recall rates for all kits, especially Agilent v8. All kits achieved an F-measure above 95.87%. Nanodigmbio had the highest precision with the fewest false positives but a slightly lower F-measure than other kits. This study also highlights the performance of new Chinese solutions from Vazyme and Nanodigmbio, which were comparable to Agilent v8 and Roche KAPA kits. These findings assist researchers and clinicians in selecting appropriate exome capture solutions.

## Introduction

Whole exome sequencing (WES) is an essential technique in genomics, focusing on the protein-coding regions of the genome. Over the past decade, the integration of WES into routine clinical diagnostics has increased due to its ability to enhance diagnostic yield. It has considerably improved the identification of genetic variants linked to diseases, making it invaluable for research and clinical applications (1,2,3,4). However, despite becoming a first-tier genetic test (5,6), the laboratory procedure remains highly complicated and sensitive without unified standards or guidelines. Achieving comprehensive coverage and uniformity, ensuring high sensitivity and specificity, and maintaining cost-effectiveness remain critical challenges in WES technology. Consequently, new comparisons of exome enrichment kits are published annually as new versions emerge (7-12).

Typically, comparison methods involve evaluating panel design, metrics of enrichment quality such as the percentage of target region coverage with the same amount of data, GC bias and the quality of variant calling in terms of sensitivity and specificity. Researchers also introduced new quality assessment metrics, such as genotypability, which measures the accuracy of read mapping (10); Cohort Coverage Sparseness and Unevenness Scores for detailed assessments of read coverage distribution (13); capture efficiency for indicating effective data usage (9) etc. All of these parameters are directly influenced by the probe design of each manufacturer, including whether they use RNA or DNA probes. Also importantly, even slight changes in the composition of hybridization and wash buffers can significantly change the efficiency of hybridization and subsequent capture procedures, potentially leading to the dropout of functionally important genes (14,15,16). It has been previously identified that approximately 1Mb of the human exome can be skipped during sequencing (17,18). Recent research has precisely localized and described these difficult-to-sequence regions in exome data, which are mainly affected by low-mappability regions, such as pseudogenes, tandem repeats, homopolymers, and other low-complexity regions (19). While WGS increases the diagnostic yield over WES (20), understanding WES’s challenges could drive improvements, minimize limitations and make it a more cost-effective procedure.

Historically, kits provided by leading companies like Agilent (SureSelect Human All Exon kits), Roche (NimbleGen SeqCap kits), and Illumina (TruSeq and Exome kits) have been widely used. Recently, high-performance kits from companies like Twist and KAPA HyperExome have emerged. Comparative studies on the Twist exome capture kit, latest Agilent v8 exome kit highlight advancements in coverage uniformity and cost-efficiency, showcasing the evolution of WES technology (7). Additionally, new competitors from China, such as Vazyme and Nanodigmbio, have introduced their own versions of exome enrichment kits with DNA probes at lower prices, further diversifying the market. In this study, we conducted a comparative analysis of four exome enrichment kits, including Agilent SureSelect Human All Exon v8, Roche KAPA HyperExome Probes, Vazyme VAHTS Target Capture Core Exome Panel and Nanodigmbio NEXome Plus Panel v1. For benchmarking, we utilized our laboratory genomic standard — a DNA sample whole genome sequenced multiple times with well-characterized variants in VCF file. This study aims to provide insights into the performance and reliability of these available to date kits.

## Material and methods

### Ethics Statement

This study conformed to the principles of the Declaration of Helsinki. The appropriate institutional review board approval for this study was obtained from the Ethics Committee at the Pirogov Medical University. All patients provided written informed consent for sample collection, subsequent analysis, and publication thereof.

### Sample Preparation and Sequencing

To conduct a comparative analysis, we enriched two pools, each containing 12 DNA libraries (24 patient DNA samples in total), using four different protocols and reagent sets: Agilent SureSelect Human All Exon v8 probes, Vazyme VAHTS Target Capture Core Exome Panel, Roche KAPA HyperExome Probes, and Nanodigmbio NEXome Plus Panel v1. DNA libraries for enrichment from the 24 initial DNA samples were prepared using the MGI Universal DNA Library Prep Set under consistent fragmentation conditions on a Covaris sonicator, followed by size selection to obtain fragments with a peak length of 250 bp. The concentrations of the prepared libraries were measured using the Qubit Flex (Life Technologies) with the dsDNA HS Assay Kit. Quality control of the DNA libraries was performed using the High Sensitivity DNA assay with the 2100 Bioanalyzer System (Agilent Technologies).

Enrichment of pools with Agilent probes was performed according to the RSMU_exome protocol (21), while the enrichment of pools with Roche, Vazyme, and Nanodigmbio probes followed the manufacturers’ hybridization kits and protocols. Subsequently, a total of 8 enriched library pools were circularized and sequenced in paired-end mode on the DNBSEQ-G400 platform using the DNBSEQ-G400RS High-throughput Sequencing Set PE100 kit, according to the manufacturer’s protocol (MGI Tech), achieving an average coverage depth of 100x. FastQ files were generated using the manufacturer’s software, zebracallV2 (MGI Tech).

### Bioinformatics pipeline

The quality of the obtained 96 paired fastq files was analysed using FastQC v0.11.9 (22). Based on the quality metrics, the fastq files were trimmed using BBDuk by BBMap v38.96 (23). Reads were aligned to the indexed reference genome GRCh38.p14 using bwa-mem2 v2.2.1 (24). SAM files were converted into BAM files and sorted using SAMtools v1.9 to check the percentage of the aligned reads (25). To correctly estimate the enrichment and sequencing quality, all 96 exomes were downsampled to 50 million reads using Picard DownsampleSam v2.22.4 (26). Based on the obtained BAM files, the quality metrics of exome enrichment and sequencing were calculated using Picard v2.22.4, and the number of duplicates was calculated using Picard MarkDuplicates v2.22.4. We performed the quality control analysis with the following bed files: Agilent v8_regions, Nanodigmbio, Roche, Vazyme. Bed files for the GENCODE and RefSeq databases were uploaded from the UCSC Table Browser (https://genome.ucsc.edu/cgi-bin/hgTables). Variant calling was performed using bcftools mpileup v1.9 (27) and DeepVariant v1.5.0 (28). After variant calling, variants in vcf files were normalized using vt normalize v0.5772 (29).

### Statistical analysis

Statistical tests were performed by Python3. To determine if the variables are normally distributed, the Shapiro–Wilk test was used. If we didn’t reject H0 hypothesis, ANOVA was performed. Otherwise, the Kruskal-Wallis test was used. Pairwise comparisons were performed for significantly different distributions. For normally distributed variables Tukey’s test was used. Otherwise, Dunn’s test was used. P-value < 0.05 as the level of statistical significance was used. Some libraries with extreme AT dropout values were excluded from the comparisons, as we consider this to be an error during the library preparation step.

## Results

### Comparison of probe designs

Contemporary platforms have reduced their target sizes to approximately 30 Mb, focusing exclusively on exonic regions. For instance, the BED file generated using Gencode V44 includes coordinates for exons from 228,189 regions, totaling 39.63 Mb (gencode_hg38.bed). Historically, manufacturers’ target sizes ranged from 50 to 60 Mb, whereas the current enrichment kits have smaller ones: Agilent V8 Exome at 35.13 Mb, Vazyme at 34.13 Mb, Nanodigmbio at 35.17 Mb, and Roche KAPA at 35.55 Mb. Significantly, 92.14% (33.86 Mb) of these regions are common across all four kits (Fig. 1). The intersections of BED files from each enrichment kit with Gencode V44 database were as follows: 86.76% (34.73 Mb) for Agilent V8, 83.80% (33.63 Mb) for Vazyme, 83.74% (34.09 Mb) for Nanodigmbio, and 84.85% (34.51 Mb) for Roche KAPA; with RefSeq database: 82.12% (34.39 Mb) for Agilent V8, 81.35% (33.76 Mb) for Vazyme, 81.43% (34.25 Mb) for Nanodigmbio, and 80.76% (34.26 Mb) for Roche KAPA. The substantial proportion of target regions, specifically 80.74% (33.53 Mb) for Gencode and 77.73% (33.61 Mb) for Refseq, are common across all four kits and databases. These results underscore the consistency and overlap in exome target size among these platforms.

**Figure 1.**
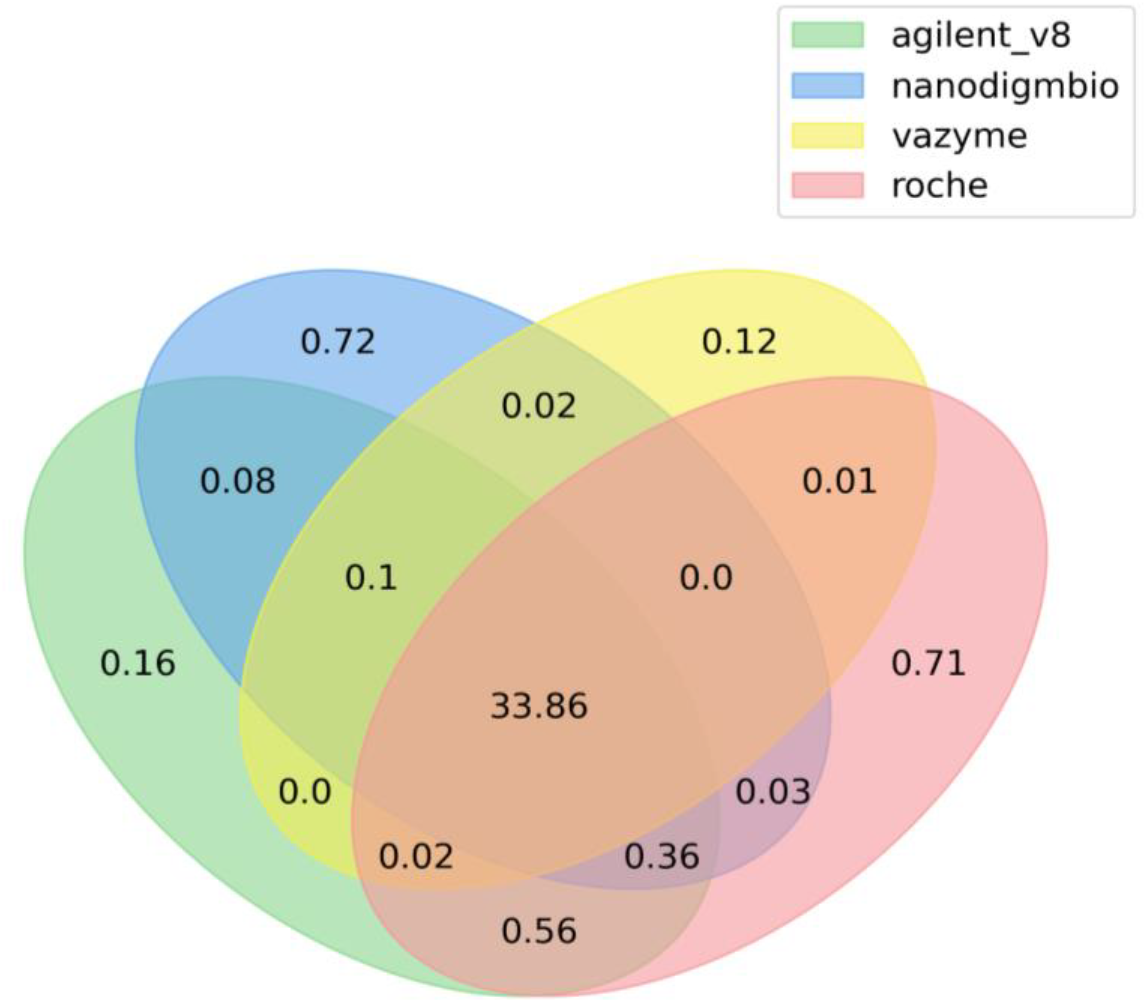
Venn diagram for intersection of Agilent SureSelect Human All Exon v8, Roche KAPA HyperExome Probes, Vazyme VAHTS Target Capture Core Exome Panel and Nanodigmbio NEXome Plus Panel v1 BED files with regard to their target sizes in megabases.

### Target coverage comparison

The mean number of reads generated for each exome kit across 96 paired fastq files (24 samples enriched by each set of probes, split into two pools) was 69,5 M, with a range of 49 to 97 M. There were no issues with mapping reads to the hg38 reference, with over 99.99% of reads aligned for all kits. To assess the enrichment quality, raw reads were downsampled to 50 M reads (one sample with 49 M reads was excluded from analysis), and the coverage statistics were obtained using Picard, with the metrics averaged across the samples. These results are detailed in Supplementary Table 1 for each downsampled pool. The mean numbers of on-target, off-target reads, and the percentage of duplicates were then determined for each platform (Fig. 2). While the number of duplicates and off-target reads varied among platforms, the mean number of on-target reads was similar for Agilent V8, Roche, and Vazyme platforms (76-78%), with Nanodigmbio showing a higher number (87%) due to fewer off-target reads. This reduction in off-target reads can be seen in IGV when comparing specific regions across all four kits in the same sample (Fig. 3). The results of the statistical tests conducted on key comparison metrics are provided in the supplementary stable 1.

**Table 1.**
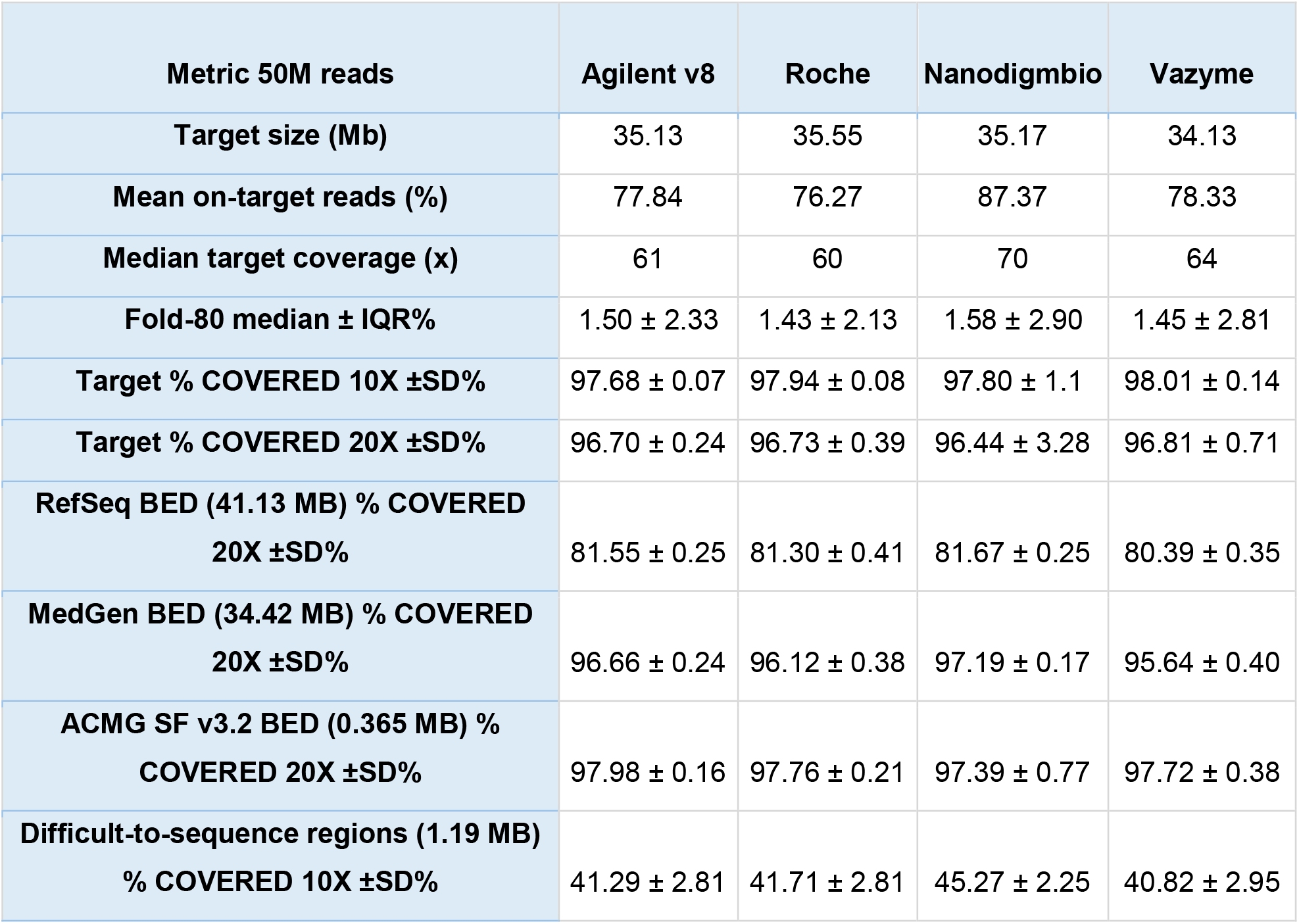
Enrichment statistics for 50M samples across four WES kits.

**Figure 2.**
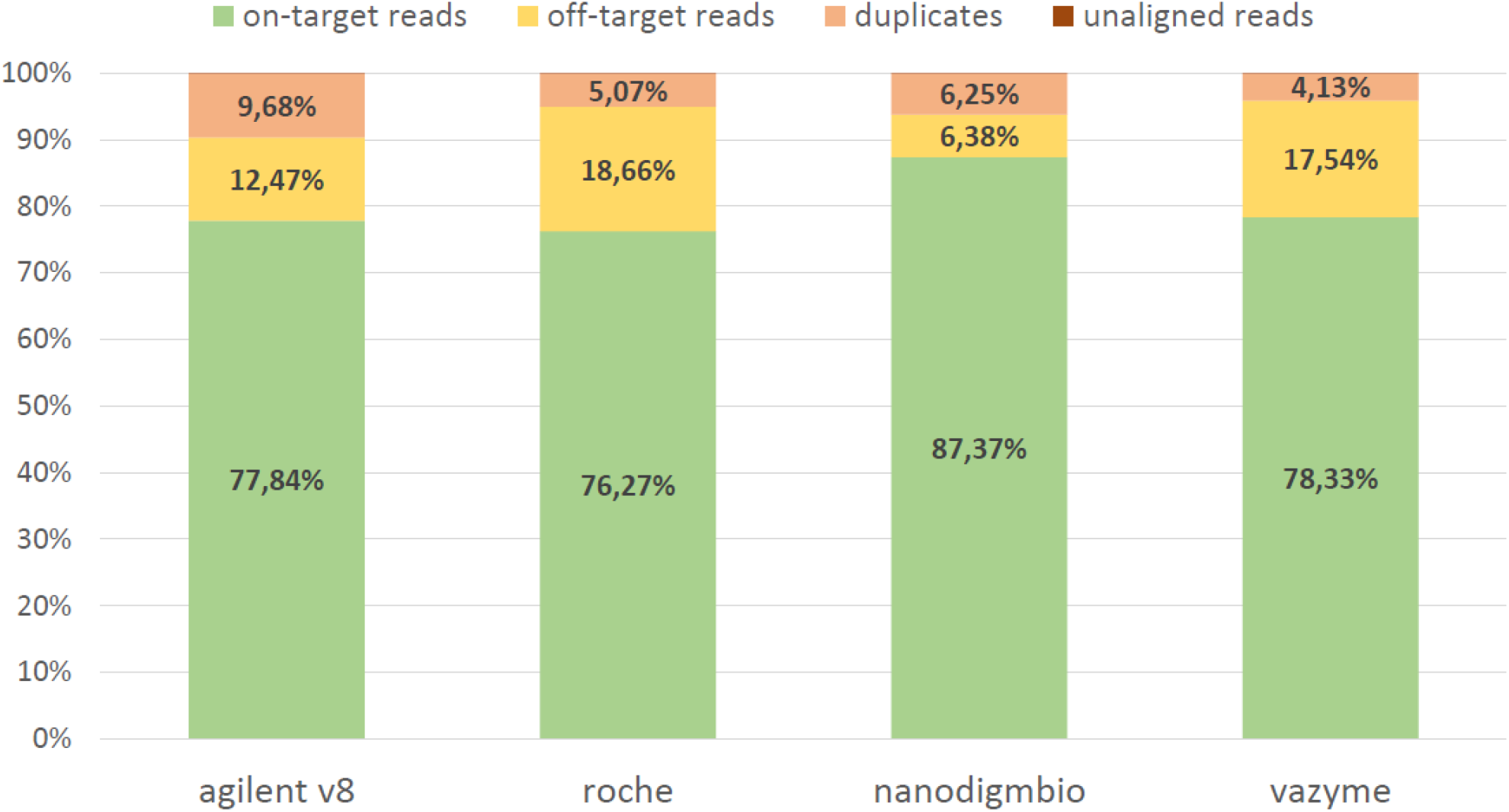
The stacked barplots for averaged on-target, off-target, duplicates, and un-aligned reads values in the pools.

**Figure 3.**
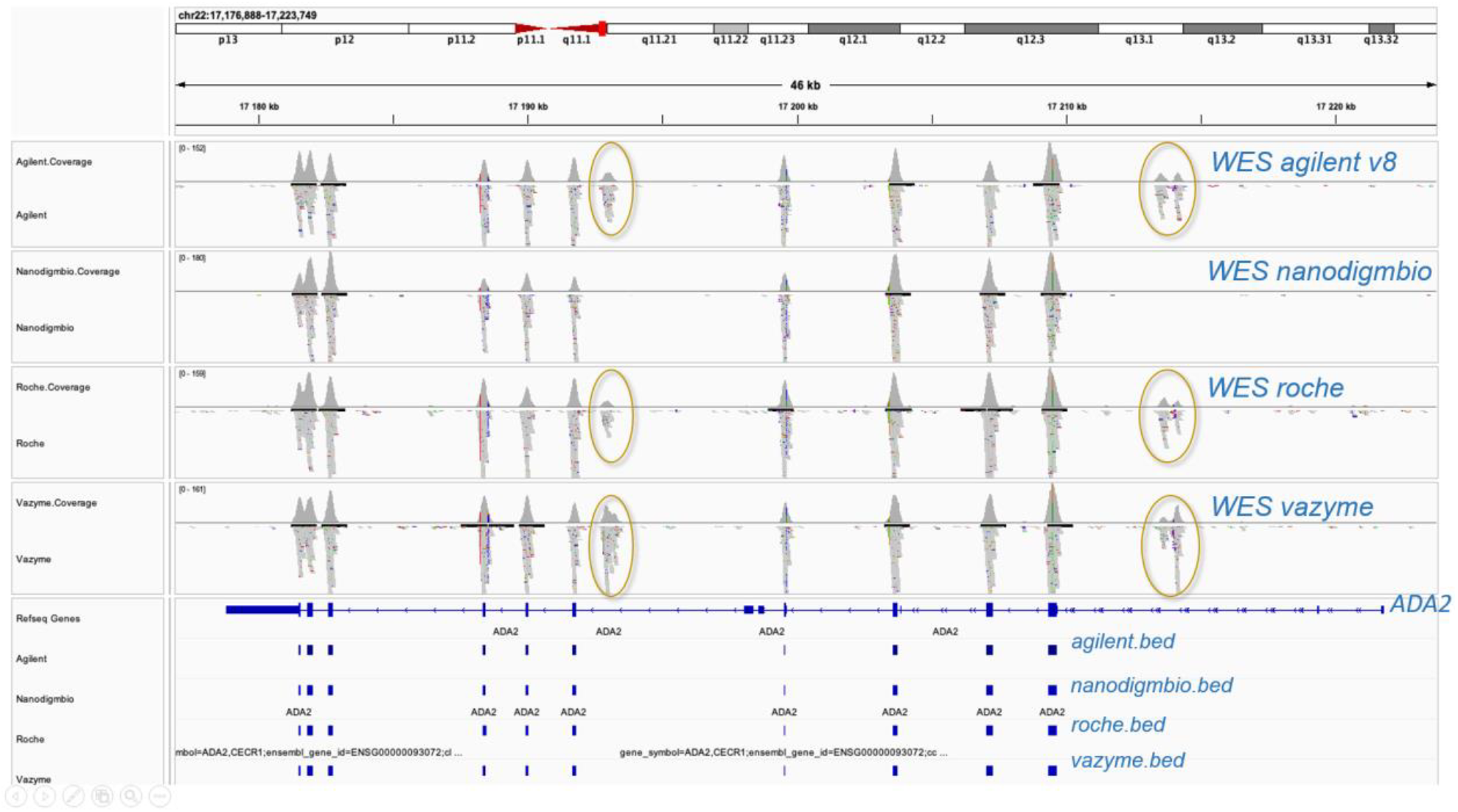
This figure shows the coverage of genomic regions of ADA2 gene on chromosome 22 across WES kits for the same DNA. Blue tracks below represent the target regions from BED files. Off-target reads are highlighted with circles, showing that same off-target reads are reproducible across Agilent, Vazyme and Roche kits but not with Nanodigmbio.

Median target coverage was slightly different for the four kits, with values of 61x and 60x for Agilent V8 and Roche KAPA, and 64x and 70x for Vazyme and Nanodigmbio respectively (Table 1). Chinese solutions showed a slight increase in median target coverage.

Regarding the fold-80 metric, which reflects the uniformity of coverage, lower fold-80 scores indicate higher uniformity, meaning all target bases are sequenced with similar coverage. Uniform coverage reduces the amount of sequencing required to reach sufficient depth for all regions of interest. The fold-80 median ± IQR% values were as follows: Agilent V8 at 1.50 ± 2.33%, Roche at 1.43 ± 2.13%, Vazyme at 1.45 ± 2.81%, and Nanodigmbio at 1.58 ± 2.90%. Despite higher median coverage for Chinese solutions, Roche exhibited the lowest fold-80 score, suggesting more uniform coverage.

Next, we evaluated the classical metric of exome enrichment – the breadth of coverage – which represents the proportion of the target covered at least x times given a specified number of reads per sample. At 50 million reads, the mean±SD breadth of coverage at 10x exceeded 97.5% for all solutions: Agilent V8 at 97.68 ± 0.07%, Roche at 97.94 ± 0.08%, Vazyme at 98.01 ± 0.14%, and Nanodigmbio at 97.80 ± 1.1%. Additionally, the breadth of coverage at 20x was above 95% for all solutions: Agilent V8 at 96.70 ± 0.24%, Roche at 96.73 ± 0.39%, Vazyme at 96.81 ± 0.71%, and Nanodigmbio at 96.44 ± 3.28% (Table 1, Figure 4a). Furthermore, we performed sequential downsampling of samples in increments of 10 million reads and found that at approximately 30 million reads, % of target covered x10 reached 95% for all sets (Figure 4b, Supp. table 1).

**Figure 4.**
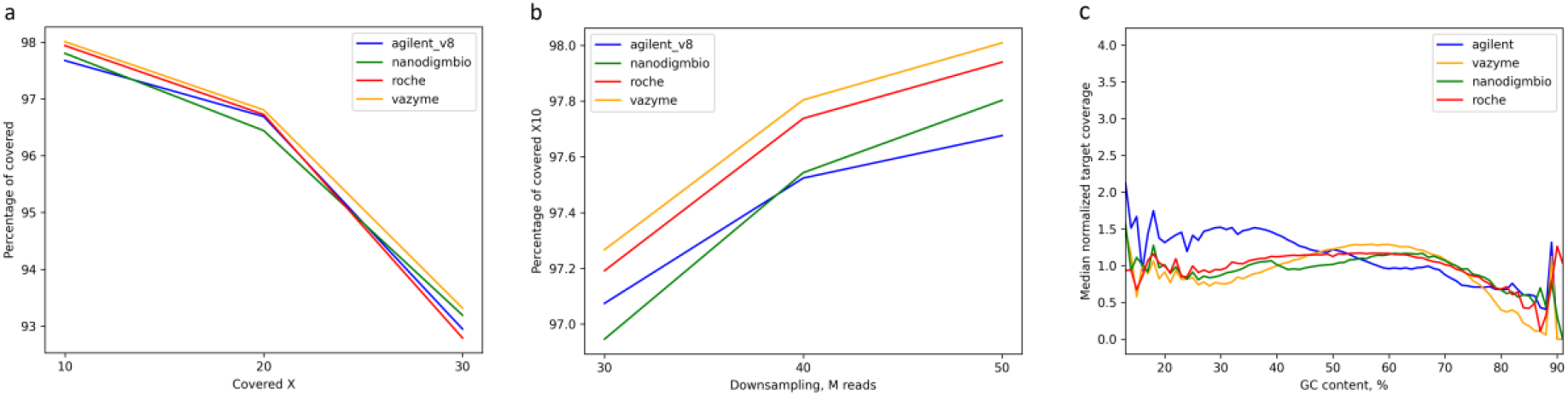
The performance of exome protocol in terms of coverage quality on downsampled samples: a) Dependence of breadth of target coverage for different depths on 50 Mb read samples; b) The percentage of regions with × 10 coverage for downsampled samples (30M, 40M, 50M) and corresponding bed files; c) GC-bias plot. The x-axis represents the GC content percentage calculated for each target region using bedtools nuc, while the y-axis shows the normalized read coverage.

In addition to standard BED sets, we assessed the breadth of the obtained exomes for the RefSeq database BED file (41.13 MB). We did not perform calculations for the GENCODE BED file because the latest versions of GENCODE (V44, V43, V42) include alternative regions (e.g., NT_187580.1) that are not present in the reference genome GRCh38. For the RefSeq BED file, the average x10 coverage values were similar between the sets and ranged from 81.14% to 82.68%, with median coverage values as follows: Agilent V8 at 55x, Roche at 55x, Vazyme at 58x, and Nanodigmbio at 62x. Also, we note that such differences in the coverage of the RefSeq BED with the authors (7) may be due to the initial different filtering of the BED file.

### GC content

To study the GC bias effect, we analyzed the normalized coverage (specific region coverage/median coverage) at different GC content of target regions for the four probe sets (Agilent v8, Vazyme, Nanodigmbio, Roche). The plot in Fig. 4c shows normalized coverage versus GC content. The Agilent platform, which uses RNA baits, exhibits the highest normalized coverage across a broad range of GC content and, as expected, shows particularly strong performance at <30%GC. The Vazyme, Nanodigmbio, and Roche platforms, all using DNA baits, demonstrate similar performance across mid-range GC content (30% to 70%) and low GC-content <30% but Vazyme tends to have a lower coverage at high GC content (>70%) than other platforms.

### Coverage comparison of difficult to sequence regions and ACMG genes

The recent advancements by Hijikata et al. (19), who have thoroughly described and compiled a list of difficult-to-sequence regions (1.19 Mb in sum) in exome datasets have enabled us to analyze how each set performed in covering these regions. Evaluating depth metrics is not meaningful since the median coverage for none of the samples exceeded 5x (Supp. table 2). Unfortunately none of the sets performed extraordinary better than the others, with the average x10 breadth as follows: Agilent V8 at 41.29 ± 1.16%, Roche at 41.71 ± 1.17%, Vazyme at 40.82 ± 1.20%, and Nanodigmbio at 45.27 ± 1.02%. These regions are characterized by low mappability and the presence of segmental duplications, tandem repeats and homopolymers, bad promoters, a %GC content <25% or >65%, etc. These factors present significant challenges under current laboratory protocols and short-read sequencing technologies.

**Table 2.**
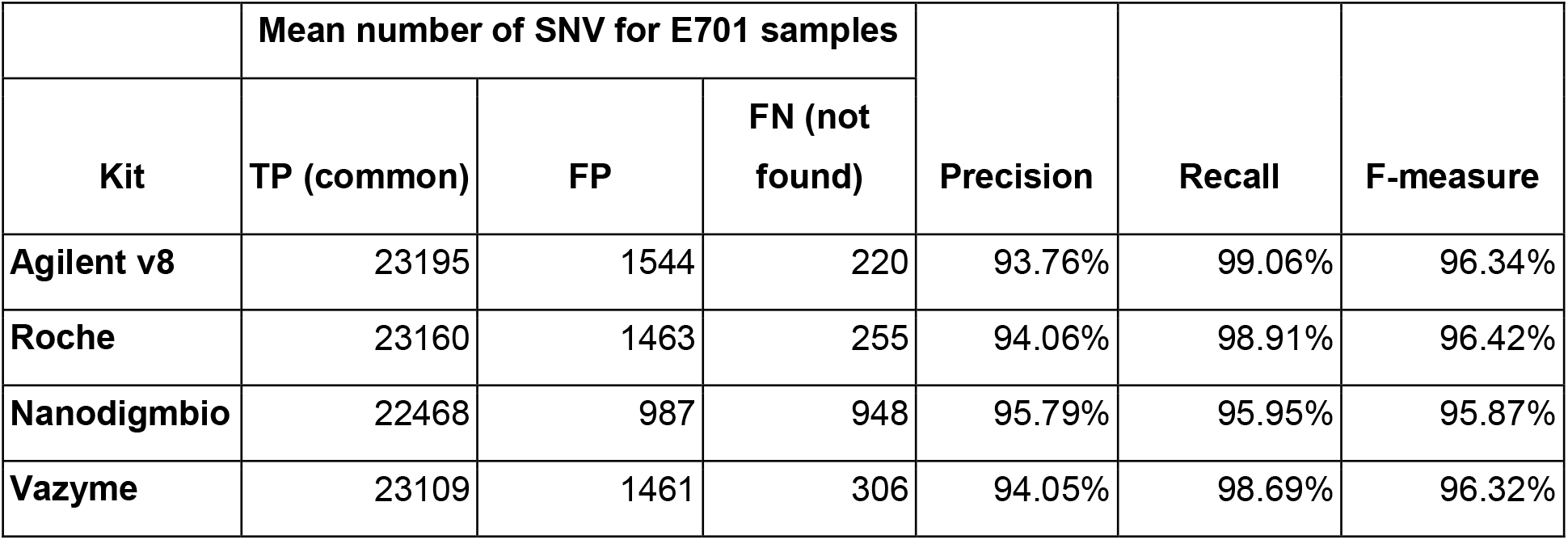
Comparative Analysis of SNV Detection Accuracy for E701 DNA standard sample. True Positive (TP) - common SNVs present in both VCF files (standart WGS E701 and experimental WES E701). False Positive (FP) - SNVs found only in the experimental E701 VCF file. False Negative (FN) - SNVs found only in the standard WGS E701 VCF file and not found in experimental E701 VCF.

We also utilized the MedGen database to compile a list of genes associated with diseases. MedGen, accessible through NCBI, provides a file containing gene IDs, MedGen entries, and OMIM IDs. These data were matched with the genemap2.txt file from OMIM.org/downloads to retrieve coordinates of disease-associated OMIM entries. Using these coordinates, we intersected them with a BED file containing coding sequences (CDS) for hg38. At 50 M reads, the mean ±SD percentage of the target regions with 20 × coverage for the MedGen BED (34,42 Mb) was as follows: Agilent V8 at 96.66±0.24%, Roche at 96.12±0.38%, Vazyme at 95.64±0.40%, and Nanodigmbio at 97.19±0.17%.

Further, we composed a Bed file (365 Kb) for 81 genes recommended by the American College of Medical Genetics and Genomics (ACMG) for pathogenic variant detection and clinical reporting (ACMG SF v3.2) (30) and calculated its coverage across the kits (Table 1, Supp. table 3). The average % of target covered x10 exceeded 98.2% for all kits. When calculating the regions from our BED ACMG file that were not included in the enrichment sets, regions with overlapping coordinates were filtered out. The total number of missing regions was: 65 regions for Agilent v8 (0.22%), 82 regions for Vazyme (0.28%), 78 regions for Nanodigmbio (0.27%), and 70 regions for Roche (0.31%) (Supp. table 3).

Among these regions were specific exons of genes, such as MUTYH, BTN, BRCA1. Additionally, some regions with coverage ranging from ∼0-3x, included in our BED ACMG file according to the UCSC browser, contained introns that flank the exons from various mRNA transcripts, such as those of the PCSK9 and MSH2 genes. In contrast, low-covered regions (from ∼3x and above 10x) were more likely to be part of the target but were difficult-to-sequence, such as exons of the TTN (Fig. 5) and PMS2 genes, which overlapped with segmental duplications and CNVs.

**Figure 5.**
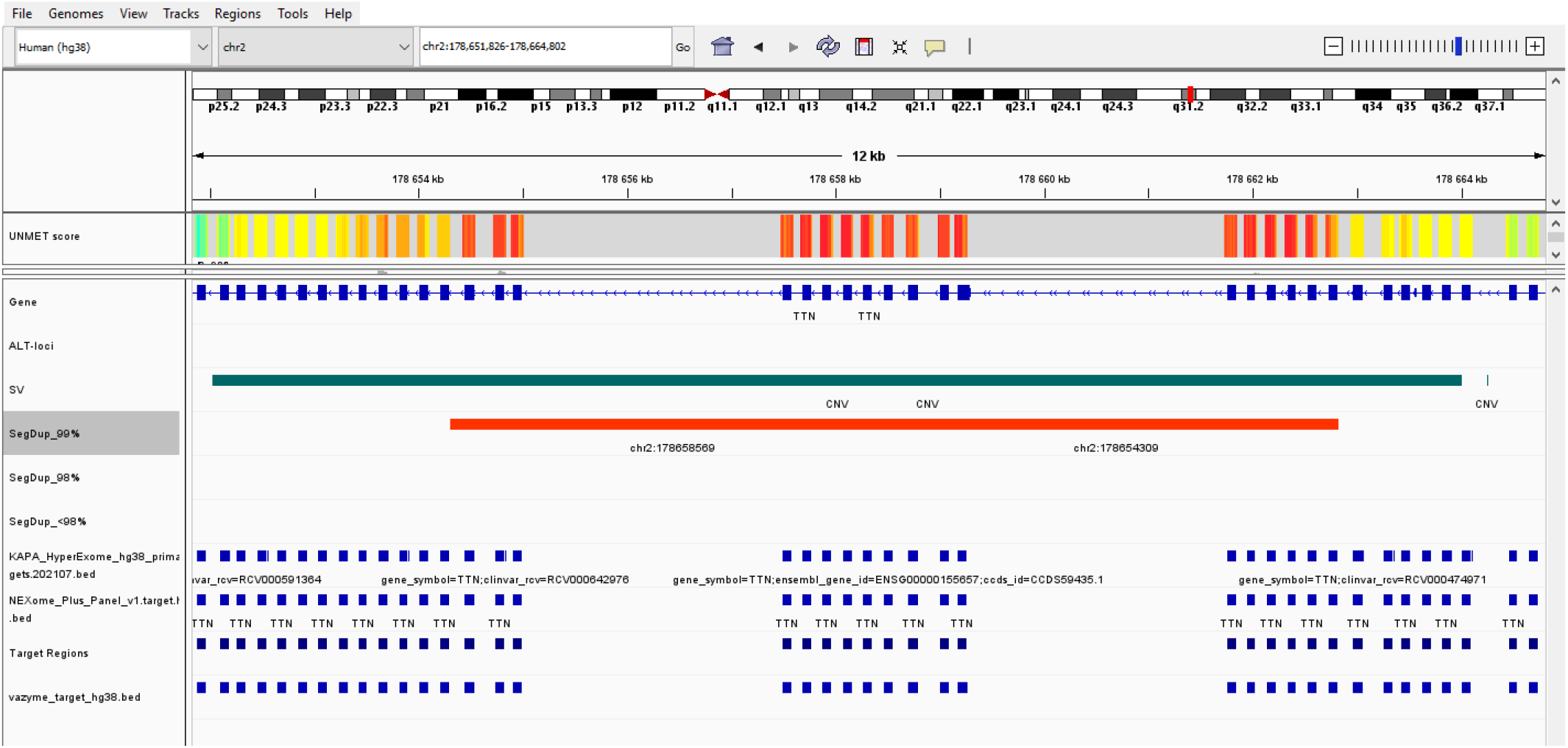
Example showing the visualization of TTN gene on IGV. Blue tracks below are exons included in target regions of BED files from different WES kits. The upper tracks show the presence of segmental duplication (SegDup) and CNV that overlap target regions, making TTN gene difficult to sequence. The UNMET score indicates the degree of difficulty in sequencing these regions (19).

### Variant calling with standard DNA

Next, we compared the ability of different exome capture kits to accurately identify genetic variants, specifically single nucleotide variants (SNVs) and small insertions and deletions (indels). Filters applied during variant calling included DP ≥13 for both bcftools and DeepVariant callers, and FILTER=PASS exclusively for DeepVariant. The number of variant counts, averaged for each enrichment kit with samples step-by-step downsampled per 10 million reads, are presented in Figure 6. The graph shows that the number of called SNPs begins to plateau at approximately 30M reads for each kit. However, this trend is not observed for indels, where the number of variants continues to increase with more reads. Interestingly, despite having a smaller target size, the highest number of variants is found in samples enriched with the Nanodigmbio kit, whereas typically more variants appear in kits with larger target size.

**Figure 6.**
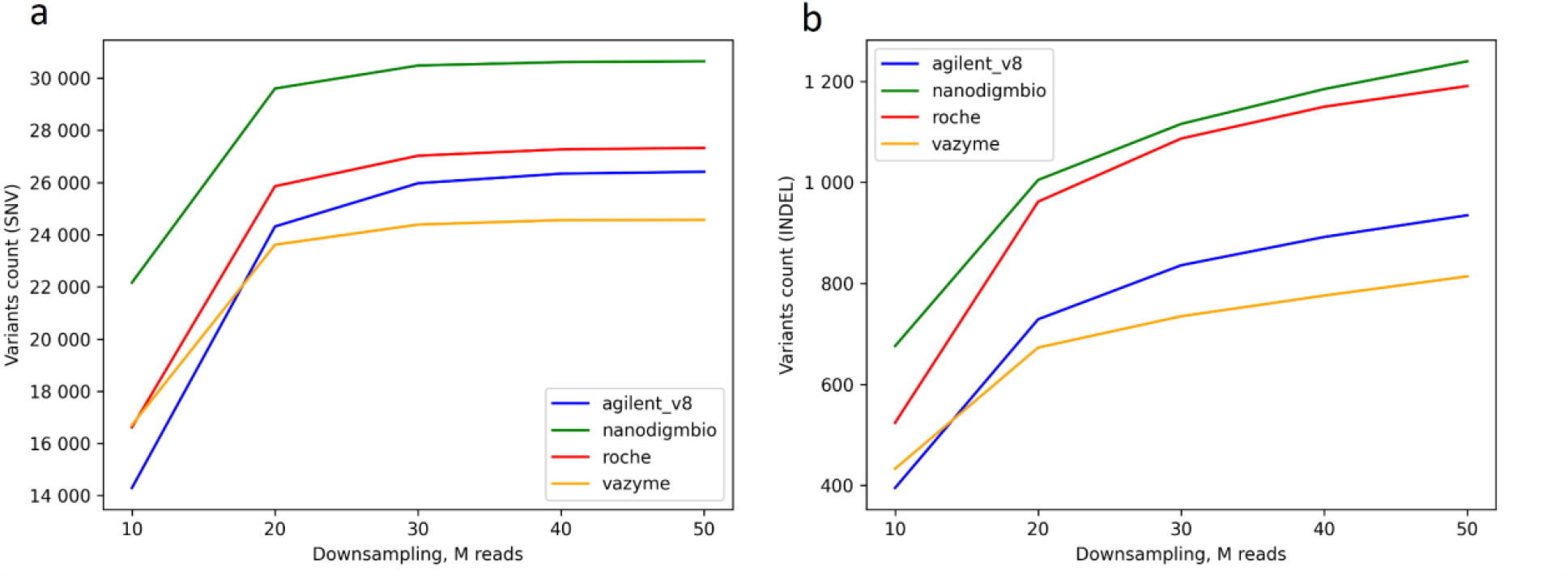
Variant calling by different WES kits (averaged for E701 samples in each pool) as a function of increasing read counts for a) SNVs, b) indels.

To benchmark the variant calling results, we used our standard human DNA sample E701. For this, we compared the exomes of the E701 standard sample (a total of 6 per each enrichment type) to the well-characterized VCF file of E701 WGS (available at: https://e701.ru). We created a BED_intersect file with a common target for all four enrichment kits and filtered VCF files for both exomes and genomes of sample E701 accordingly. Then experimental E701 WES VCF files were intersected with E701 WGS using bcftools isec, and precision, recall, and F-measure were calculated (Table 2, Supp. table 4).

The Nanodigmbio kit showed the most distinct results from the other kits, with the highest number of not found variants and the lowest number of false positive (FP) variants. The other kits showed similar results with a slight increase in FP variants and fewer “not found” variants. The recall rates were exceptionally high for all kits, particularly for Agilent V8, indicating a strong ability to identify most true variants present in the sample. The F-measure, which balances precision and recall, was similar for Roche, Agilent V8, and Vazyme, all above 96%. Nanodigmbio had a slightly lower F-measure, though still robust at 95.87%.

## Discussion

All four kits demonstrated strong performance metrics, making them reliable choices for exome sequencing. Among them, Roche provided the most uniform exome coverage. However, Roche, Agilent V8 and Vazyme may be preferable for applications where maximizing recall is essential, while Nanodigmbio is recommended for scenarios where precision is paramount. This study is the first to evaluate the new Chinese solutions from Vazyme and Nanodigmbio, which, in our opinion, perform comparably to the established Agilent V8 and Roche KAPA kits. We aimed to use our in-house genomic standard — a DNA sample that has been sequenced at least three times using whole genome sequencing. This approach allowed us to assess the variant calling performance of each kit, rather than drawing conclusions based solely on coverage data.

The fact that fold-80 values could vary for the same kit for the same sample types, but in different laboratories, shows the multifactorial nature of the enrichment process and its sensitivity to many laboratory techniques and protocols such as DNA extraction, library preparation kits, DNA insert size, etc. Additionally, differences in results between studies may be due to various approaches to the bioinformatic processing of exome data and the lack of a standardized “golden pipeline” (31).

When compared to previous studies from our laboratory (8,21), which also used downsampling to 50 million reads, it is evident that modern kits show less steep declines in key metrics such as fold-80 and breadth of coverage. This improvement is due to advances in probe design and hybridisation protocols. For example, new for the market kits - Roche KAPA, Vazyme, Nanodigmbio - show similar performance to previously described last Agilent v8 kit in maintaining high % of target coverage even at lower read depths, highlighting the technological advances in these modern exome enrichment solutions. Previous solutions (MGI v4, Agilent v6, v7) didn’t even reach 90% of target coverage at 20x read depth, while modern solutions exceed 96%. Fold-80 metrics used to be over 2, now it’s in the 1.4-1.6 range.

Historical biases in older enrichment kits have had a significant impact on sequencing results. In the TCGA cohort of 6353 samples, the limitations of these older kits (mostly the Custom V2 Exome Bait kit) introduced significant bias. This bias resulted in a significant number of undercovered genes, leading to an undercalling of mutations in cancer genes (16). Specifically, at least 4833 genes were under-covered, highlighting the challenges of capturing exons for sequencing with earlier technologies.

Despite advances in exome capture technologies, the coverage of difficult-to-sequence regions remains a significant challenge. These regions, characterized by low mappability, segmental duplications, homopolymers, tandem repeats and extreme GC content, present significant obstacles for all exome capture kits. Unfortunately, none of the kits evaluated in this study excelled in covering these problematic areas although they didn’t show a dramatic shift in coverage of GC content. Now it becomes more clear that well-designed probes kits are less affected by GC-bias and the problem of regions with low mapping quality is more related with short read sequencing (13,17-19). We recommend that researchers consider these limitations when designing experiments, and also refer to the IGV visualization of difficult-to-sequence genes from Hijikata’s article, which we found particularly helpful. In our view, new WES solutions from Vazyme and Nanodigmbio in general offer performance on the same level as the Agilent V8 and Roche KAPA kits. Chinese kits are also recognized as being more cost-effective. Consequently, the choice of kit should take into account specific project needs and budget considerations.

## Data availability

Exome sequences will be deposited into the NCBI open-access sequence read archive (SRA) in fastq.gz format. BioProject number will be added soon.

